# Surface-based Tractography uncovers ‘What’ and ‘Where’ Pathways in Prefrontal Cortex

**DOI:** 10.1101/2024.03.21.585573

**Authors:** Marco Bedini, Emanuele Olivetti, Paolo Avesani, Daniel Baldauf

## Abstract

The frontal eye field (FEF) and the inferior frontal junction (IFJ) are prefrontal regions that mediate top-down functions, with mounting neuroimaging evidence suggesting that they specialize in controlling spatial versus non-spatial processing, respectively. We hypothesized that their unique patterns of structural connectivity underlie these specialized roles. To accurately infer the localization of FEF and IFJ in standard space, we performed an activation likelihood estimation meta-analysis of fMRI paradigms that reliably targeted these regions. Using surface-based probabilistic tractography methods at the individual subject level, we tracked streamlines ipsilaterally from the inferred FEF and IFJ activation peaks to the dorsal and ventral visual streams mapped on the native white matter surface parcellated using the multimodal Glasser atlas. By contrasting FEF and IFJ connectivity likelihoods, we found predominant structural connectivity from the FEF to regions of the dorsal visual stream compared to the IFJ (particularly in the left hemisphere), and conversely, predominant structural connectivity from the IFJ to regions of the ventral visual stream compared to the FEF bilaterally. Additionally, we analyzed the cortical terminations of the superior longitudinal fasciculus to the FEF and IFJ, implicating its first and third branches as segregated pathways mediating their communication to the posterior parietal cortex. The structural connectivity fingerprints of the FEF and IFJ support the view that the two visual stream architectures extend to the posterior lateral prefrontal cortex and provide converging anatomical evidence of their specialization in spatial versus non-spatial control.

## 1. INTRODUCTION

A longstanding tradition in modern neuroscience argues that in primates, the regions that mediate cognitive control and flexibility are primarily hosted in the prefrontal cortex (PFC; Chafee & Heilbronner, 2022; Luria, 1966; Miller & Cohen, 2001). The PFC cannot be fully understood in isolation, without considering its rich connectivity patterns (Fuster, 2001; Passingham & Lau, 2022). Indeed, several PFC regions form crucial hubs of brain networks with adaptive functions (Cole et al., 2013; Duncan, 2001; Yeo et al., 2011). In the human posterior lateral PFC, the frontal eye field (FEF) and the inferior frontal junction (IFJ) exhibit the characteristics of control regions that are involved in a wide array of functions ranging from attention, working memory, and other forms of cognitive control (reviewed in Bedini & Baldauf, 2021), and are responsible for biasing activity in posterior brain regions (Baldauf & Desimone, 2014; Gazzaley & Nobre, 2012; Veniero et al., 2021; Zanto et al., 2011; Zhang et al., 2018). Crucially, however, recent evidence suggests that these regions differ in their selectivity to spatial versus non-spatial information: on the one hand, the FEF predominantly encodes spatial information (Fiebelkorn & Kastner, 2020; Mackey et al., 2017; Sprague & Serences, 2013; Wang et al., 2015), whereas, on the other hand, the IFJ predominantly encodes non-spatial (i.e., feature-and object-based) information (Baldauf & Desimone, 2014; Bedini & Baldauf, 2021; O’Reilly, 2010). The fMRI studies that employed multivariate pattern analysis (MVPA; Peelen & Downing, 2023) to decode the attended and memorized information from the FEF and IFJ provide evidence supporting this hypothesis (discussed in Bedini & Baldauf, 2021). The dissociation in the selectivity of neural populations of these lateral prefrontal regions is in line with the model that was originally proposed by Goldman-Rakic and colleagues to understand the organization of the PFC in non-human primates based on neurophysiological and tract-tracing evidence (Constantinidis & Qi, 2018; Goldman-Rakic, 1996). In particular, tracer evidence suggests that the dorsolateral and ventrolateral PFC were characterized by divergent patterns of anatomical connectivity to the ‘what’ and ‘where’ visual pathways (Goodale & Milner, 1992; Mishkin et al., 1983), respectively (Felleman & Van Essen, 1991; Romanski, 2004; Wilson et al., 1993; Yeterian et al., 2012). In humans, anatomical connections are primarily inferred using diffusion MRI (dMRI) tractography (Jbabdi et al., 2015; Jeurissen et al., 2019; Van Essen et al., 2014), but to date, the distinct structural connectivity patterns of the regions in the posterior lateral PFC remain elusive.

Previous dMRI studies were able to reveal that many aspects of the organization of prefrontal connectivity are preserved across the macaque and the human brain (Jbabdi et al., 2013a; Neubert et al., 2014; Sallet et al., 2013; Thiebaut de Schotten et al., 2012), which allows to formulate hypotheses motivated by comparative evidence (Mars et al., 2021). Other influential studies on the PFC focused on performing a connectivity-based parcellation to infer areal borders based on tractography (Thiebaut de Schotten et al., 2017; Tomassini et al., 2007). This technique hinges on the idea that a characteristic pattern of connectivity (i.e., the connectivity fingerprint; Passingham et al., 2002) defines each brain region (Eickhoff et al., 2018), which in turn fundamentally underlies its specialized role in cognition (Mars et al., 2018). Each PFC region’s unique set of afferent and efferent connections ultimately constrain the inputs it receives (and consequently its selectivity for specific sensory information), as well as its ability to control and bias activity elsewhere in the brain to solve particular tasks, respectively (Passingham & Lau, 2022; Petrides et al., 2012). Diffusion MRI studies combining the functional localization of regions of interest at the individual subject level using fMRI, with the analysis of their structural connectivity with tractography, offered initial evidence on the involvement of long-range association pathways in mediating the functions associated with the FEF and IFJ (reviewed in Bedini & Baldauf, 2021). Of importance to the present study, the studies by Anderson et al. (2012), Sczepanszki et al. (2013), and Umarova et al. (2010) provided tractography evidence of FEF wiring patterns to the posterior parietal cortex. Baldauf and Desimone (2014) showed that the connectivity likelihood with the fusiform face area and the parahippocampal place area increased in the ventrolateral PFC and near the IFJ. Collectively, these studies were important milestones in suggesting that these regions communicate with the parietal and temporal cortices through distinct, reciprocal, and potentially segregated white matter pathways. These putative pathways are known to be supported by specific white matter association bundles (Barrett et al., 2020; Thiebaut de Schotten et al., 2012) that can be delineated using virtual dissection techniques via dMRI tractography (Catani & Thiebaut de Schotten, 2008; Warrington et al., 2020). A seminal dMRI study indicated that a major white matter bundle that mediates the communication within the dorsal and ventral attention networks (Corbetta & Shulman, 2002; Yeo et al., 2011) is the superior longitudinal fasciculus (SLF), which is subdivided into three branches that run from the dorsomedial to the ventrolateral direction (Thiebaut de Schotten et al., 2011). Damage to the SLF often leads to visual neglect, especially in the case of right hemisphere damage (Bartolomeo & Seidel Malkinson, 2019; Doricchi et al., 2008). In addition to lesion evidence, the functions of the SLF have also been inferred by combining tractography with meta-analytic techniques. According to a large-scale study combining the virtual dissection of the SLF in 129 participants with 14 fMRI meta-analyses, the SLF1 primarily mediates ‘spatial-motor’, and the SLF3 ‘non-spatial-motor’ functional components (Parlatini et al., 2017). In a recent comprehensive study, Sani et al. (2021) analyzed dMRI data from 263 participants released by the Human Connectome Project (HCP; Van Essen et al., 2013a) to understand the organization of the networks involved in attention control. The authors found reliable tractography evidence that the human posterior inferotemporal area (phPIT) forms part of an interconnected network encompassing the FEF and the lateral intraparietal area dorsal (LIPd), two well-known areas forming the core of the dorsal attention network. The authors then attempted to uncover the white matter bundles underlying these results. Interestingly, they found that the connectivity between the FEF and LIPd is mediated by the SLF2 and in part by the SLF3, and the connectivity between the FEF and phPIT is mediated by the posterior arcuate fasciculus and the inferior fronto-occipital fasciculus, although there was only moderate overlap with the former bundle.

Overall, the studies discussed above offer some evidence of the structural connectivity patterns of posterior lateral PFC regions (mainly the FEF, and in a more limited way, the IFJ) and the underlying white matter bundles (SLF), and help clarify the function of these connections within the attention networks. However, these studies also faced a tradeoff between having relatively small sample sizes but with the great advantage of having regions of interest (ROIs) functionally localized at the individual level, as opposed to the ability to analyze large datasets but with regions defined at the group level or by using an atlas, which may not always represent the individual brain organization faithfully (Eickhoff et al., 2018). Even though the functional localization of ROIs at the individual level seems to be the most accurate way to overcome the issues associated with group-level analyses performed on brain templates (Coalson et al., 2018; Fedorenko, 2021), the localization of the ROIs may be impractical to scale to larger samples. As a consequence of the former limitation, another major barrier in adequately mapping the connectivity fingerprints of a brain region and relating them to its functions is the fact that most of the available brain atlases aren’t tailored to incorporate information from large collections of task-based fMRI data (Eickhoff et al., 2018). Therefore, in the present study, we first performed an activation likelihood estimation (ALE; Eickhoff et al., 2012) fMRI meta-analysis to overcome these issues and accurately infer the localization of FEF and IFJ in standard space. We used two samples of studies using functional localizers and other well-replicated fMRI paradigms that target these regions (as in Bedini et al. 2023), which enabled us to investigate their structural connectivity in a large dataset, such as the HCP (Van Essen et al., 2013a). Although the studies briefly reviewed here suggest that the FEF and the IFJ may communicate with posterior parietal and temporal regions through distinct and segregated pathways (Anderson et al., 2012; Baldauf & Desimone, 2014; Thiebaut de Schotten et al., 2011; Sani et al., 2021; Sczepanszki et al., 2013; Umarova et al., 2010), no study has yet directly contrasted their structural connectivity to the posterior visual regions. A divergence in these connectivity patterns would be consistent with the principles of anatomical connectivity discovered in the macaque PFC (Romanski, 2004; Yeterian et al., 2012), in which case we predict that the FEF should be more likely connected to the dorsal stream, and the IFJ to the ventral stream. Such results would highlight how long-range white matter pathways underlie the functional specialization of these regions in spatial versus non-spatial control processes. Therefore in the present study, we sought to investigate the structural connectivity fingerprints of the FEF and IFJ with posterior visual regions. We combined an ALE meta-analysis - which allowed us to accurately infer the localization of these regions in standard space - with a surface-based probabilistic tractography analysis at the individual subject level - to contrast their connectivity likelihoods to the global dorsal and ventral visual streams. Finally, we also performed the virtual dissection of the major association bundles connecting the PFC and posterior regions (Barrett et al., 2020; Catani & Thiebaut de Schotten, 2008) to uncover the white matter pathways underlying the hypothesized connectivity patterns. We specifically focused on the cortical terminations of the SLF, as we predicted that its three branches would have distinct patterns of projections to the FEF and IFJ.

## 2. MATERIALS AND METHODS

### 2.1 Subjects and Data sources

We used the MRI data from the HCP-MEG subjects release (Van Essen et al., 2013b) available at: https://db.humanconnectome.org. Our sample included 56 unrelated subjects (age: 22-35, mean = 28.66 ± 3.81; see the Supplementary Information for the selection criteria) and its size is considerably larger compared to some of the previous dMRI studies cited. This factor should enable more reliable inferences with dMRI tractography, where the research community is still involved in large-scale efforts to validate the various models and algorithms available (Maier-Hein et al., 2017; Sarwar et al., 2021; Sotiropoulos & Zalesky, 2017).

### 2.2 Localization of the FEF and IFJ seed regions

We performed an ALE fMRI meta-analysis to accurately infer the localization of the FEF and IFJ in the MNI152 space using GingerALE (v. 3.0.2; Eickhoff et al., 2012). This analysis was carried out on part of the experiments that were included in our previous meta-analysis (Bedini et al. 2023; see Supplementary Information, Tables S1 and S2 for details), which we later refined and expanded in that study. The ALE parameters were set to *p* = 0.01, with a cluster-level family-wise error (FWE) = 0.01 and 1000 permutations.

### 2.3 Seeding approach

The FEF and IFJ ALE peak coordinates in the MNI152 space were non-linearly mapped using FSL FNIRT to the native dMRI space (Figure 1A). These peaks were then projected onto the WM/GM boundary of each subject’s (the low-resolution native white matter surface, i.e. with 32k vertices) available in the HCP extended structural preprocessing folder (containing FreeSurfer recon-all segmentation outputs) as follows. We first created a sphere of a 2.5mm radius centered around the ALE peak. Each FEF and IFJ seed therefore consisted of a 6.25mm diameter sphere at this stage. We note that by this method, our spheres included the coordinates corresponding to the ALE peaks reported in Bedini et al. (2023). Next, we used the command-line tool surf_proj from FSL to map these spheres to the WM/GM boundary from the FreeSurfer segmentation (i.e., *h_white.gii). The latter file was mapped from the FreeSurfer space to the diffusion space using the FSL surf2surf command. We used the ‘mean’ method in surf_proj, and we further increased the step size by the voxel size (i.e., 1.25mm) until each subject had identifiable vertices corresponding to the volumetric spheres (see Figure 1A for examples). The final step size was set to 2.5mm for every subject. Initializing the streamlines from seeds projected onto the WM/GM boundary has the advantage of mitigating some tractography biases (Sotiropoulos & Zalesky et al., 2017), including most importantly the gyral bias (Girard et al., 2014; Schilling et al., 2018; Van Essen et al., 2014).

**FIGURE 1.**
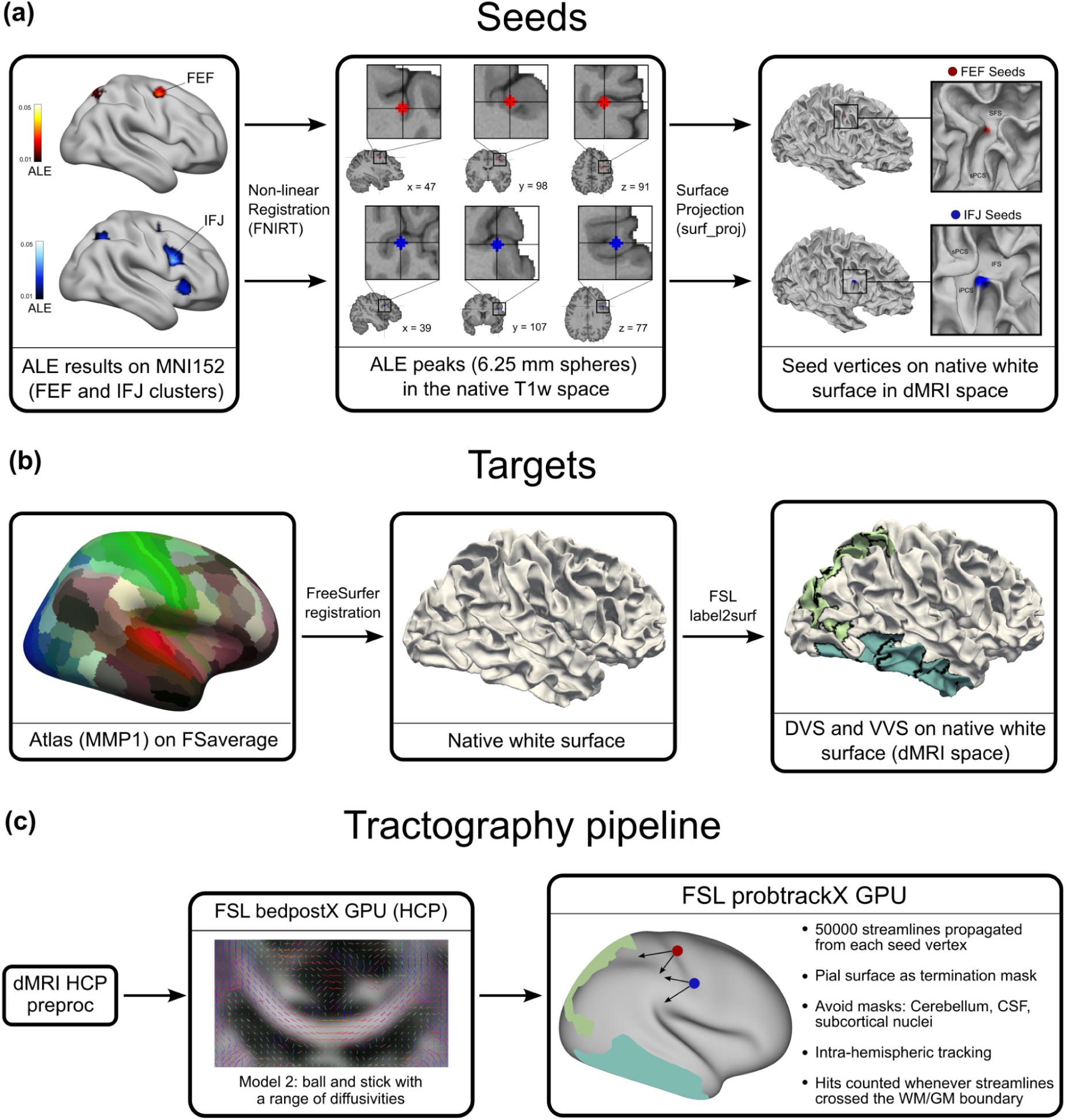
A. ALE results and projection of the FEF and IFJ peaks to the WM/GM boundary of an example subject. **B** Mapping of the dorsal and ventral visual stream regions from the MMP1 atlas to the native white surface of the same example subject. **C** Overview of the surface-based tractography pipeline. From the preprocessed data, the fiber orientations were modeled by HCP’s bedpostx, and we propagated streamlines ipsilaterally from each seed vertices (FEF, IFJ) in the individual subject space and counted hits on the target visual streams using FSL probtrackx GPU (Hernandez-Fernandez et al., 2019)

### 2.4 Atlas selection and definition of the target regions of the two visual streams

Recent approaches to parcellate the human cortex increasingly leverage multiple sources of information from different modalities (Eickhoff et al., 2018). The multimodal parcellation (MMP1) published by Glasser et al. (2016) provides a comprehensive understanding of the organization of the human cortex combining information about architecture, function, retinotopy, and connectivity, allowing us to systematically investigate the connectivity of the FEF and IFJ to the dorsal and ventral visual streams defined using these multimodal criteria. The MMP1 was projected on Fsaverage (Fischl et al., 1999) by Kathryn Mills following the indications from Coalson et al. (2018) and is available at: https://figshare.com/articles/dataset/HCP-MMP1_0_projected_on_fsaverage/3498446. The resulting FreeSurfer annotation files from the MMP1 atlas were mapped to the native structural space using the tool developed by the CJ Neurolab (https://figshare.com/articles/dataset/HCP-MMP1_0_volumetric_NIfTI_masks_in_native_str <u>uctural_space/4249400</u>; FreeSurfer v6.0, Fischl, 2012; see Figure 1B). We highlight that surface-based registration methods achieve much better consistency in labeling cortical areas and in capturing intersubject variability compared to traditional volumetric methods (Coalson et al., 2018). Therefore, by using the procedure described above to map the MMP1 parcellation to the native dMRI space, we could retain high fidelity in mapping this atlas to the individual cortical anatomy. After these steps, we registered all the labels from the atlas onto the white matter/grey matter (WM/GM) boundary of each subject’s (the low-resolution white matter surface, i.e., with 32k vertices) as provided by the FreeSurfer segmentation (v6.0; Fischl, 2012) using the FSL command label2surf. To select the parcels of the MMP1 atlas that belonged to the dorsal and ventral visual streams, we consulted comparative (Kravitz et al., 2011, 2013; Mishkin et al., 1983) and human (Goodale & Milner, 1992) evidence for their definition, guided by the accompanying materials from the MMP1 publication (see Glasser et al., 2016, supplementary information #3). In addition, we took into account another defining characteristic of the dorsal visual stream, namely the presence of topographic visual organization (Wang et al., 2015), to better refine the inclusion criteria for the parcels belonging to this stream (see Table 1, and the Supplementary Information for details). We also compared our definitions to a previous study that investigated the connectivity of the extrastriate body area with these streams as defined by the MMP1 (Zimmermann et al., 2018). In contrast to that study, since we were not concerned about confounds due to the excessive proximity of the targets to the seed regions, we included additional parcels from the MMP1 that were discarded by the authors for that reason (Zimmermann et al., 2018).

**TABLE 1.** List of the MMP1 parcels belonging to the dorsal and ventral visual streams.

| Parcel | Stream | Full name | Parcel | Stream | Full name |
| --- | --- | --- | --- | --- | --- |
| V3A | Dorsal | Area V3A | V8 | Ventral | Eight Visual Area |
| V3B | Dorsal | Area V3B | VVC | Ventral | Ventral Visual Complex |
| V6 | Dorsal | Sixth Visual Area | PIT | Ventral | Posterior InferoTemporal |
| V6A | Dorsal | Area V6A | FFC | Ventral | Fusiform Face Complex |
| V7 | Dorsal | Seventh Visual Area | VMV1 | Ventral | VentroMedial Visual Area 1 |
| IPS1 | Dorsal | IntraParietal | VMV2 | Ventral | VentroMedial Visual Area 2 |
| 7AL | Dorsal | Lateral Area 7A | VMV3 | Ventral | VentroMedial Visual Area 3 |
| 7PL | Dorsal | Lateral Area 7P | TE1a | Ventral | Area TE1 anterior |
| 7PC | Dorsal | Area 7PC | TE1m | Ventral | Area TE1 middle |
| LIPd | Dorsal | Area Lateral IntraParietal dorsal | TE1p | Ventral | Area TE1 posterior |
| LIPv | Dorsal | Area Lateral IntraParietal ventral | TE2a | Ventral | Area TE2 anterior |
| VIP | Dorsal | Ventral IntraParietal Complex | TE2p | Ventral | Area TE2 posterior |
| MIP | Dorsal | Medial IntraParietal Area | TF | Ventral | Area TF |
| IP1 | Dorsal | Area IntraParietal 1 | PHT | Ventral | Area PHT |
| V3CD | Dorsal | Area V3CD | PH | Ventral | Area PH |
| LO3 | Dorsal | Area Lateral Occipital 3 | TGv | Ventral | Area TG ventral |
| MT | Dorsal | Middle Temporal Area |  |  |  |

### 2.5 Probabilistic tractography method

We performed all the analyses of dMRI data using FSL (v. 6.0.5; Jenkinson et al., 2012). Susceptibility-induced distortions, eddy currents, and subject motion were corrected using the HCP preprocessing pipelines (Andersson & Sotiropoulos, 2016; Van Essen et al. 2013b). All the dMRI data was analyzed in the subjects’ native space and the original voxel resolution (i.e., 1.25mm). First, we ran FSL DTIFIT on the preprocessed data to obtain the fractional anisotropy image, which we used for performing image registrations to the native dMRI space. To infer the connectivity likelihoods between the FEF and IFJ seeds and the target regions, we implemented the FSL FDT analysis pipeline, which involves two steps, namely bedpostx (Jbabdi et al., 2012) and probtrackx (Behrens et al., 2007). Since the first step is computationally intensive and given the advantages offered by the GPU implementation of bedpostx (Sotiropolous et al., 2016), we downloaded the outputs of bedpostx released by the HCP (<u>HCPpipelines/bedpostx</u>) and used them as starting points for our probabilistic tractography pipeline. The following sections describe in detail our methodology to run probtrackx.

### 2.6 Surface-based probabilistic tractography with GPU acceleration

Probtrackx samples the fiber orientation estimated with bedpostx to generate a connectivity distribution between user-defined seed and target regions (Behrens et al., 2007). Importantly, probtrackx estimates how robust the connection is against noise and uncertainty, or in other words, how reproducible it is (Jbabdi & Johansen-Berg, 2011). This method allowed us to infer how likely a region is on the dominant pathway of anatomical connectivity from a seed region (Jeurissen et al., 2019). We leveraged the GPU implementation of probtrackx on a CUDA 9.2 parallel architecture, which achieves massive speedups compared to the CPU version (Hernandez-Fernandez et al., 2019). To perform this step, we split the list of targets into four sub-groups due to the exceeding memory demands of the computations. Overall, our general approach to probabilistic tractography closely follows the study by Donahue et al. (2016) and more specifically, their method for generating the dense connectome matrix (termed dDT1 in that study). We chose to follow this tracking approach as the authors showed that it improved the agreement between measures of connectivity derived from probtrackx and retrograde tracing in the macaque, with a much higher correlation factor than previous studies. We set probtrackx parameters as follows: 50000 streamlines were propagated from each seed vertex for a maximum of 3200 steps using modified Euler integration. The curvature threshold was set to 0.2 (corresponding to an 80° turn) with a step length of 0.3125 mm (1/4 of the voxel size), and loopcheck was enabled to discard artifactual loops of the streamlines. Similarly to Donahue et al. (2016), we also tracked the streamlines ipsilaterally, enabling us to add several anatomical constraints to tractography. We used the corpus callosum, subcortical nuclei, cerebellum, and cerebrospinal fluid masks resulting from the FreeSurfer segmentation as ‘avoid’ masks. The native pial surface was instead used as a ‘termination’ mask since this approach is effective in preventing invalid jumps of the streamlines between adjacent gyri (Hernandez-Fernandez et al., 2019). Whenever a streamline crossed the WM/GM boundary, the streamline count was increased for the cortical target corresponding to that triangular mesh. Our approach builds on the growing dMRI literature that focuses on the problem of assigning streamline terminations to cortical areas (Schilling et al., 2018; St-Onge et al., 2018; Yeh et al., 2019), and the evidence suggesting that surface-based tractography provides clear advantages when handling more complex geometrical configurations that can’t be resolved using conventional volumetric techniques (Cottaar et al., 2021; Shastin et al., 2022; St-Onge et al., 2021; Yeh et al., 2019). For the control analysis focusing on the distance confounding factor, we repeated the same probtrackx command as previously described and enabled the ‘pd’ flag, which multiplies the streamline count for each parcel by the average distance traveled by the respective streamlines from the seed.

### 2.7 Post-processing, statistical analysis, and data visualization

Because our seeding strategy involved the propagation of the streamlines from a variable number of vertices across subjects, the total number of streamlines initialized is given by 50000 times the number of vertices in each seed. The individual raw streamline counts were summed across vertices within each label, and we computed the ratio between the surviving streamline count and the total number of streamlines propagated from each seed region that were not discarded based on the avoid masks (termed waytotal in probtrackx) to obtain a normalized streamline count (NSC)^2^ for each subject. Following that, we averaged the NSC across subjects for each seed to target region pairs. We then contrasted each seed (FEF, IFJ) to target (dorsal, ventral stream) NSC performing paired *t*-tests with *p* = 0.01 and correcting for multiple comparisons using the false discovery rate (FDR; Benjamini & Hochberg, 1995) with *q* < 0.05. We also performed a control analysis to account for a potential distance confound by repeating the same statistical contrasts on the distance-corrected probtrackx outputs. We performed the statistical analyses using JASP (https://jasp-stats.org), and we computed the FDR-adjusted p-values using the web utility provided by the software Seed-based d Mapping (https://www.sdmproject.com/utilities). The visualizations of the cortical connectivity values were created using Connectome Workbench (v. 1.5.0; Marcus et al., 2013). The RGB values corresponding to the connectivity likelihoods were generated using custom code written in MATLAB2019b (https://www.mathworks.com) based on the diverging colormaps developed by Brewer (https://colorbrewer2.org). Finally, the raincloud plots (Allen et al., 2021) were created by modifying the Python example code from: https://github.com/pog87/PtitPrince. All the NSCs were log_2_-scaled for visualization convenience since we tracked streamlines from a variable number of vertices across subjects, which led to non-linear increases in the surviving streamline count.

### 2.8 Virtual dissection of the association white matter bundles, analysis of their asymmetries, and the cortical terminations of the SLF

Warrington et al. (2020) developed a method for the automatic virtual dissection of the major human white matter bundles in the HCP dataset. In summary, this method leverages predefined waypoints, exclusion, and termination masks, and initializing probtrackx from specific seed masks enables their virtual dissection. We used this method to segment the white matter association bundles of the PFC that could potentially overlap with FEF and IFJ ALE clusters using the definitions by Barrett et al. (2020). Our bundles of interest included the three branches of the superior longitudinal fasciculus (SLF1, SLF2, and SLF3), the arcuate fasciculus (AF), and the uncinate fasciculus (UF). We used a subsample of 24 subjects randomly selected from the previous dataset. This analysis was also performed using the GPU version of probtrackx (Hernandez-Fernandez et al., 2019). We segmented all the white matter bundles of interest in the subject’s native space. The method for the analysis of the bundles’ lateralization and the associated results are reported in the Supplementary Information (p. 13). Our method to estimate the cortical projections of the three branches of the SLF involved the following steps: first, we non-linearly mapped the FEF and IFJ ALE clusters in MNI152 space to each subject’s T1w. By leveraging the subject-specific tissue segmentation, we removed all the voxels of the clusters that were found within the white matter. We then dilated these masks using a 3×3×3 mm kernel. This step was crucial to ensure that the clusters reached the white matter and followed closely previous approaches to estimating white matter bundles’ terminations (e.g., Pestilli et al., 2014). To quantify the likelihood of cortical terminations to each ALE cluster, we normalized the streamline count by the number of overlapping voxels between each SLF branch and the FEF and IFJ ALE clusters separately. We contrasted the streamline count projecting to the FEF and IFJ clusters for the SLF1, SLF2, and SLF3 by performing *t*-tests with *p* = 0.01.

## 3. RESULTS

### 3.1 Surface-based probabilistic tractography results

We contrasted the NSC for each seed to the ipsilateral targets separately in the left and right hemispheres (LH, RH) using Wilcoxon’s signed-rank test (*w*) since the data violated the normality assumption (Shapiro-Wilk test). The *p* values reported were corrected for multiple comparisons using the FDR (*q* < 0.05). The contrast between the LH FEF and the LH IFJ NSC revealed that in the dorsal visual stream, several regions had a higher connectivity likelihood with the LH FEF compared to the LH IFJ (see Figure 2A; in the following, we always list the parcels from the most anterior to posterior): 7PC (*w*, *p* < 0.001, *r_B_* = 0.722), 7AL (*w*, *p* < 0.001, *r_B_* = 0.9), LIPd (*w*, *p* < 0.001, *r_B_* = 0.65), LIPv (*w*, *p* < 0.001, *r_B_* = 0.706), VIP (*w*, *p* < 0.001, *r_B_* = 0.934), IP1 (*w*, *p* < 0.001, *r_B_* = 0.843), MIP (*w*, *p* < 0.001, *r_B_* = 0.872), 7PL (*w*, *p* < 0.001, *r_B_* = 0.875), IPS1 (*w*, *p* < 0.001, *r_B_* = 0.924), V7 (*w*, *p* = 0.001, *r_B_* = 0.648), V6A (*w*, *p* < 0.001, *r_B_* = 0.703), V3CD (*w*, *p* < 0.001, *r_B_* = 0.851), and V3B (*w*, *p* < 0.001, *r_B_* = 0.945). The contrast between the LH FEF and the LH IFJ NSC revealed that in the ventral visual stream, several regions had a higher connectivity likelihood with the LH IFJ compared to the LH FEF (see Figure 2C): TGv (*w*, *p* < 0.001, *r_B_* = −0.559), TE2a (*w*, *p* < 0.001, *r_B_* = −0.657), TE1m (*w*, *p* = 0.002, *r_B_* = −0.495), and TE1p (*w*, *p* < 0.001, *r_B_* = −0.739).

**FIGURE 2.**
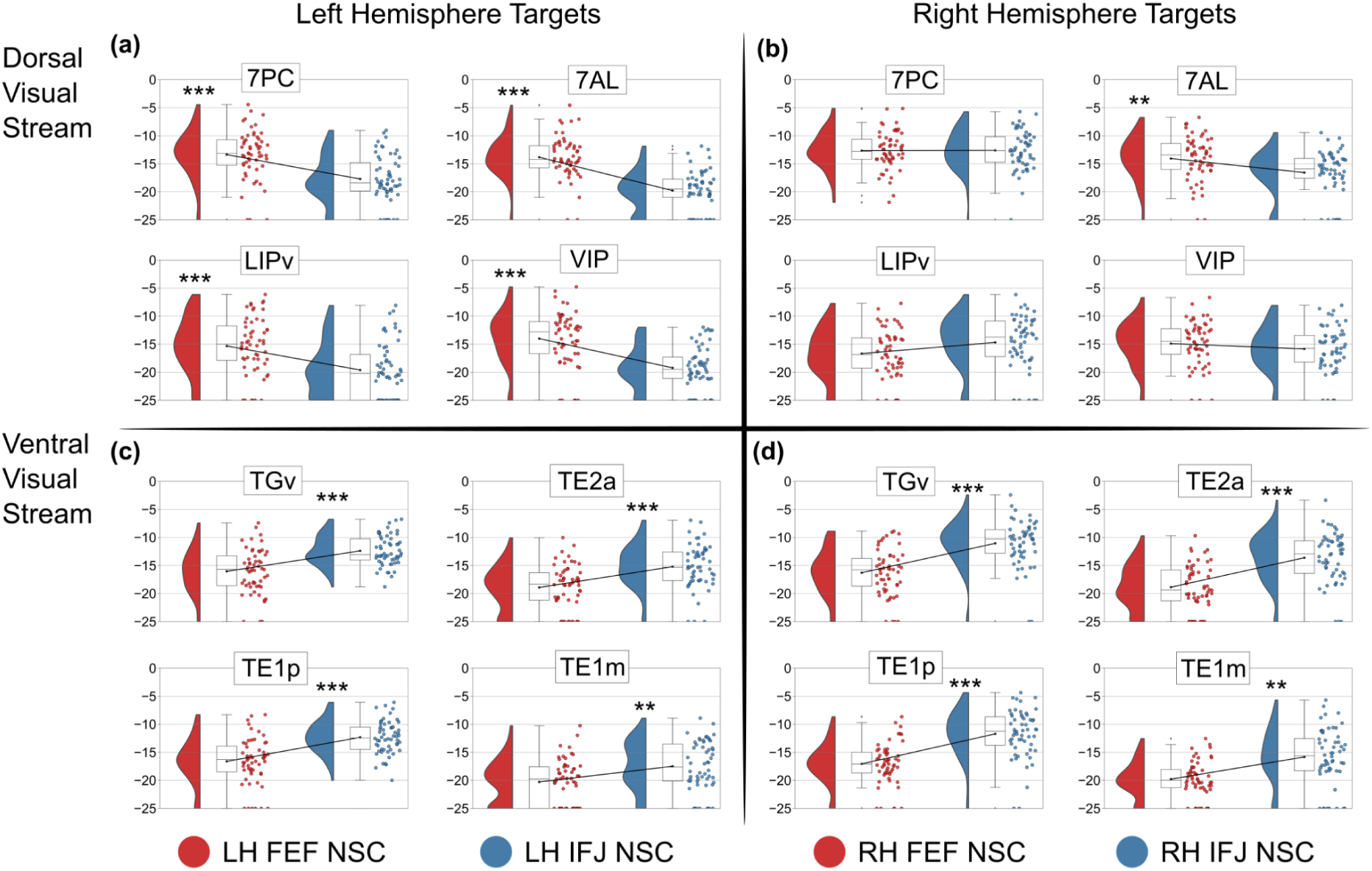
Plots of the log_2_-scaled normalized streamline counts (NSC) of the FEF and IFJ to cortical targets in the most anterior portions of the dorsal and ventral visual streams in the left (panels A and C) and right hemisphere (panels B and D). A value of −25 is set as an arbitrary log_2_ connectivity likelihood equal to 0 for visualization convenience The contrast between the LH FEF and the LH IFJ distance-corrected NSC revealed that in the ventral visual stream, several regions had a higher connectivity likelihood with the LH IFJ than with the LH FEF: TGv (*w*, *p* < 0.001, *r_B_* = −0.55), TE2a (*w*, *p* < 0.001, *r_B_* = −0.603), TE1m (*w*, *p* < 0.001, *r_B_* = −0.589) and TE1p (*w*, *p* < 0.001, *r_B_* = −0.732). The LH FEF had, in contrast, a higher connectivity likelihood with the areas VVC (*w*, *p* = 0.002, *r_B_* = 0.616), FFC (*w*, *p* = 0.005, *r_B_* = 0.486), and PIT (*w*, *p* = 0.002, *r_B_* = 0.64) than with the LH IFJ. The contrast between the RH FEF and the RH IFJ distance-corrected NSC revealed that in the dorsal visual stream, only one region had a higher connectivity likelihood with the RH FEF than with the RH IFJ, namely 7AL (*w*, *p* = 0.007, *r_B_* = 0.469) as in our main results. However, we also found that area V3A had a higher connectivity likelihood with the LH IFJ than with the LH FEF (*w*, *p* = 0.007, *r_B_* = −0.535). The contrast between the RH FEF and the RH IFJ distance-corrected NSC revealed that in the ventral visual stream, several regions had a higher connectivity likelihood with the RH IFJ than with the RH FEF: TGv (*w*, *p* < 0.001, *r_B_* = −0.906), TE2a (*w*, *p* < 0.001, *r_B_* = −0.865), TE1a (*w*, *p* < 0.001, *r_B_* = −0.712), TF (*w*, *p* < 0.001, *r_B_* = −0.602), TE1m (*w*, *p* < 0.001, *r_B_* = −0.836), TE2p (*w*, *p* < 0.001, *r_B_*= −0.729), TE1p (*w*, *p* < 0.001, *r_B_* = −0.926), FFC (*w*, *p* = 0.007, *r_B_* = −0.515), PH (*w*, *p* < 0.001, *r_B_* = −0.776), VMV1 (*w*, *p* = 0.006, *r_B_* = −0.848), and V8 (*w*, *p* = 0.007, *r_B_* = −0.723).

The LH FEF had a higher connectivity likelihood with the areas VVC (*w*, *p* = 0.006, *r_B_* = 0.550) and FFC (*w*, *p* = 0.006, *r_B_* = 0.503) compared to the LH IFJ. The contrast between the RH FEF and the RH IFJ NSC revealed that in the dorsal visual stream, only one region had a higher connectivity likelihood with the RH FEF compared to the RH IFJ (see Figure 2B), namely 7AL (*w*, *p* = 0.001, *r_B_* = 0.558). On the other hand, the contrast between the RH FEF and the RH IFJ NSC revealed that in the ventral visual stream, several regions had a higher connectivity likelihood with the RH IFJ compared to the RH FEF (see Figure 2D): TGv (*w*, *p* < 0.001, *r_B_* = −0.877), TE2a (*w*, *p* < 0.001, *r_B_* = −0.87), TE1a (*w*, *p* < 0.001, *r_B_* = −0.719), TF (*w*, *p* < 0.001, *r_B_* = −0.628), TE1m (*w*, *p* = 0.002, *r_B_* = −0.821), TE2p (*w*, *p* < 0.001, *r_B_* = −0.62), TE1p (*w*, *p* < 0.001, *r_B_* = −0.94), PH (*w*, *p* < 0.001, *r_B_* = −0.772), and VMV1 (*w*, *p* = 0.009, *r_B_*= −0.853).

To better understand our results, we investigated whether they could be due to the average streamline length between the seed and the target regions. Therefore, we reanalyzed our data using the probtrackx outputs corrected by distance. In these outputs, each surviving streamline count was multiplied by the average length traveled by the streamlines to reach their targets. The *p* values reported were also corrected for multiple comparisons using the FDR (*q* < 0.05). The contrast between the LH FEF and the LH IFJ distance-corrected NSC revealed that in the dorsal visual stream, several regions had a higher connectivity likelihood with the LH FEF than with the LH IFJ: 7PC (*w*, *p* < 0.001, *r_B_* = 0.685), 7AL (*w*, *p* < 0.001, *r_B_* = 0.901), LIPd (*w*, *p* < 0.001, *r_B_* = 0.658), LIPv (*w*, *p* < 0.001, *r_B_* = 0.64), VIP (*w*, *p* < 0.001, *r_B_* = 0.904), IP1 (*w*, *p* < 0.001, *r_B_* = 0.835), MIP (*w*, *p* < 0.001, *r_B_* = 0.857), 7PL (*w*, *p* < 0.001, *r_B_* = 0.872), IPS1 (*w*, *p* < 0.001, *r_B_*= 0.847), V7 (*w*, *p* < 0.001, *r_B_* = 0.825), V6A (*w*, *p* < 0.001, *r_B_* = 0.64), V3CD (*w*, *p* < 0.001, *r_B_* = 0.811), and V3B (*w*, *p* < 0.001, *r_B_* = 0.922).

We summarize these results by plotting the difference between LH FEF and LH IFJ, and RH FEF and RH IFJ NSCs on the Conte69 mesh (Van Essen et al., 2012) parcellated using the MMP1 (Figure 3A). We indicated all the targets with significantly higher connectivity likelihood with the FEF or IFJ in the main results and the control analysis by asterisks in Figure 3B.

**FIGURE 3.**
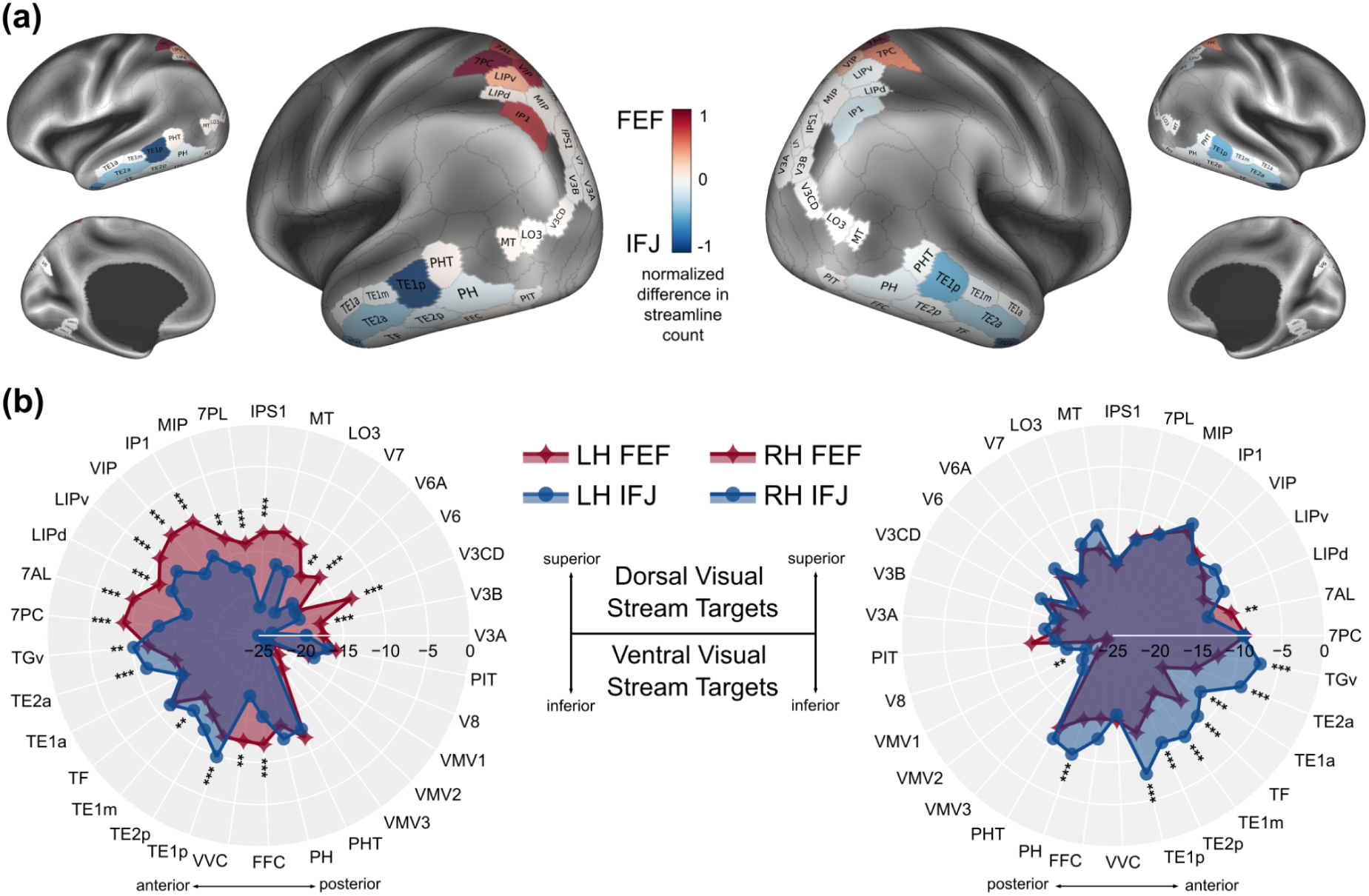
**A** Difference in the connectivity likelihood of the FEF and IFJ with the dorsal and ventral visual streams projected on the Conte69 surface parcellated using the MMP1 (Glasser et al., 2016). **B** Spider plots of the log_2_-scaled NSC of FEF and IFJ to each cortical target from the MMP1 (we set −25 as a log_2_-transformed arbitrary 0 connectivity likelihood for visualization convenience as in the previous figure). Asterisks indicate significant results that were replicated in the control analysis for distance (** = *p* < 0.01, *** = *p* < 0.001)

### 3.2 Relationship between the FEF and IFJ ALE clusters and the cortical terminations of the three branches of the SLF

By using the method we developed to quantify the cortical projections of the three branches of the SLF, we could contrast their estimated white matter terminations to the FEF and IFJ ALE clusters (expressed as a normalized streamline density count). In the left hemisphere (Figure 4A), the SLF1 had a significantly higher number of streamlines projecting to the LH FEF than to the LH IFJ (*w*, *p* < 0.001, *r_B_* = 0.98). In contrast, the SLF3 had a significantly higher number of streamlines projecting to the LH IFJ than to the LH FEF (*w*, *p* < 0.001, *r_B_* = −0.947). Similarly, in the right hemisphere (Figure 4B), the SLF1 had a significantly higher number of streamlines projecting to the RH FEF than to the RH IFJ (*w*, *p* < 0.001, *r_B_* = 0.86). In contrast, the SLF3 had a significantly higher number of streamlines projecting to the LH IFJ than to the LH FEF (*w*, *p* < 0.001, *r_B_* = −0.933). We didn’t find any significant differences in the SLF2 streamline projections in either the left or right hemisphere.

**FIGURE 4.**
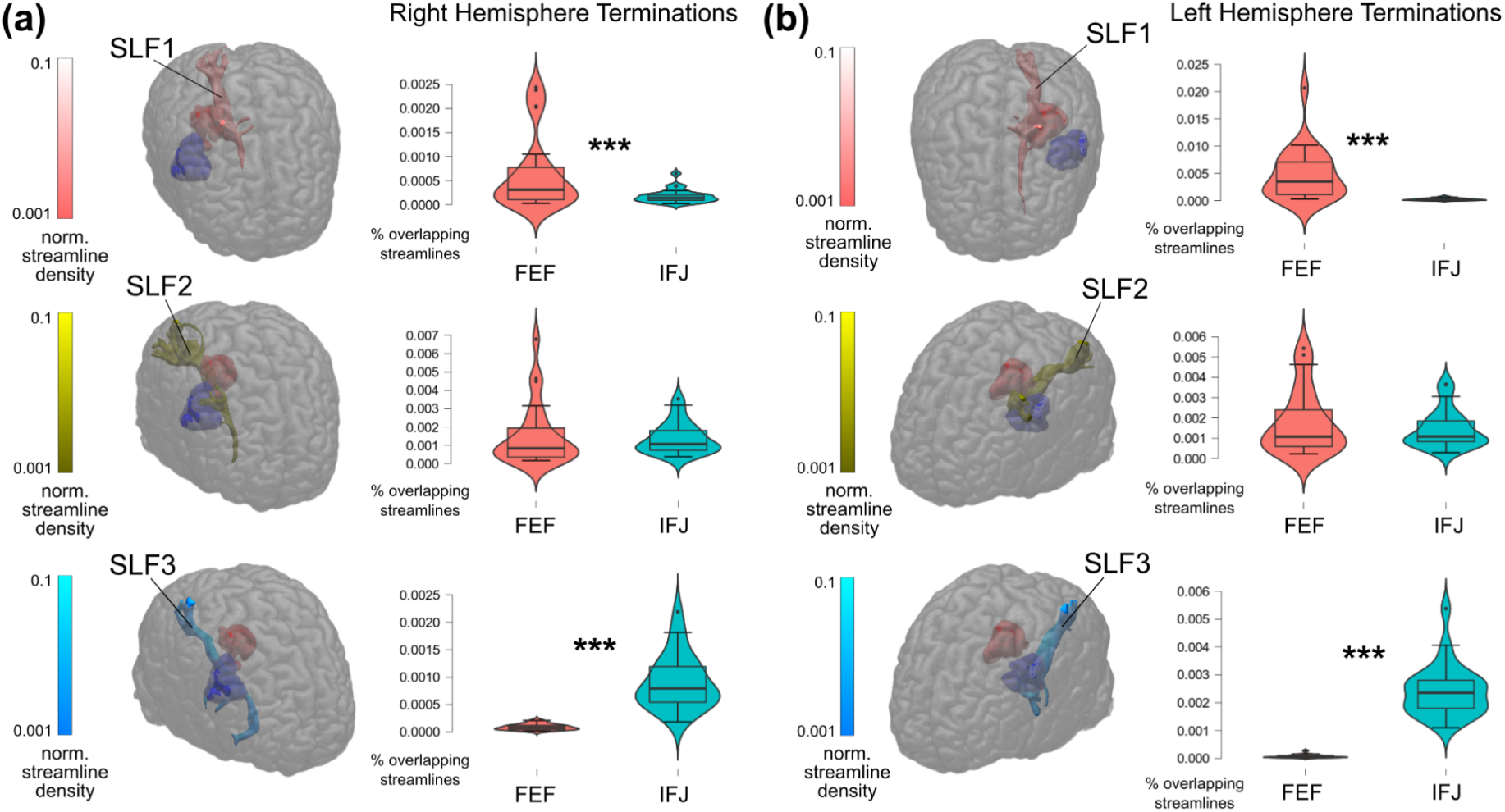
Overlay of the SLF1, SLF2, and SLF3 for an example subject with the dilated FEF (in red) and IFJ (in blue) ALE clusters, and plots of the difference in their estimated cortical terminations in the right (panel A) and left (panel B) hemisphere, respectively. In both hemispheres, SLF1 streamlines had a higher probability of projecting to the FEF ALE cluster, whereas SLF3 streamlines had a higher probability of projecting to the IFJ ALE cluster (*p* < 0.001)

## 4. DISCUSSION

The FEF and the IFJ are prefrontal regions that continue receiving interest in the field due to their involvement in several orchestrating functions (Bedini & Baldauf, 2021). Their overarching role is to provide top-down feedback to bias activity in posterior regions in visual attention and working memory tasks (Baldauf & Desimone, 2014; Gazzaley & Nobre, 2012; Veniero et al., 2021; Zanto et al., 2011; Zhang et al., 2018). Their structural connectivity patterns remained largely elusive to date, partly due to a lack of consensus on the precise localization of these regions in standard space (Bedini et al., 2023) as well as the intrinsic limitations of dMRI tractography (Jbabdi et al., 2015; Jeurissen et al., 2019; Sotiropoulos & Zalesky, 2017). In this study, we combined an ALE meta-analysis, which allowed us to accurately infer the localization of the FEF and IFJ in MNI152 space, with surface-based probabilistic tractography - to uncover their structural connectivity with the dorsal and ventral visual streams (Goodale & Milner, 1992) leveraging the high-quality dMRI data from the HCP dataset (Van Essen et al., 2013a). Our ALE results revealed the strongest convergence of activations at the junction of the superior frontal sulcus and precentral sulcus for the FEF, and at the junction of the inferior frontal sulcus and precentral sulcus for the IFJ (see Bedini et al., 2023, for an extension of these results). We parcellated each participant’s cortex using the MMP1 (Glasser et al., 2016) to define our target regions and we used spherical seeds centered on the FEF and IFJ ALE peaks to perform the surface-based tractography analysis. We propagated streamlines from the WM/GM boundary by applying multiple anatomically-motivated constraints to the tractography. By contrasting the connectivity likelihood of FEF and IFJ in each hemisphere to all the ipsilateral target regions, we found that in the left hemisphere, the FEF had predominant connectivity with several regions across the dorsal visual stream, whereas the IFJ had predominant connectivity with most regions of the ventral visual stream. In the right hemisphere, we found predominant connectivity of the FEF to only one region of the dorsal stream, and again, a predominant connectivity of the IFJ with multiple regions of the ventral stream. We generally found the strongest evidence for a divergence in these connectivity patterns in regions in the high and mid regions in the processing hierarchy of each stream, while it was almost entirely abolished in early visual regions. These patterns seem to underline a gradual rather than absolute transition in the zone of the predominance of FEF and IFJ connectivity, as these regions were likely also connected to the targets in the non-dominant stream. Another striking aspect of our results is that they suggest a remarkable hemispheric asymmetry: while in the left hemisphere, they largely supported our hypothesis in both streams, in the right hemisphere, we found results in line with our prediction primarily in the ventral stream, but not in the dorsal stream. Quantitatively, these results seem to be driven by an increased connectivity likelihood of the RH IFJ with the dorsal stream, as opposed to a decrease in the connectivity likelihood of the RH FEF per se (see Figure 3b and Supplementary Information, Figure S6). These asymmetries therefore potentially suggest a more robust involvement of the RH IFJ in the dorsal visual stream, and fit well with a recent study that examined the structural connectivity of the dorsal and ventral attention networks, showing that these networks are right-lateralized (Alves et al., 2022). We found highly consistent results in our control analysis, where we corrected the connectivity likelihoods based on the average distance traveled by the streamlines to reach their targets from the seeds, thus ruling out this potential confound. Taken together, our findings reveal a reliable anatomical motif of segregated long-range pathways connecting the FEF and IFJ to the dorsal and ventral visual streams. These results are in agreement with the tracer evidence from the macaque model (Gerbella et al., 2010; Schall et al., 1995; Stanton et al., 1993; Webster et al., 1994; Yeterian et al., 2012), and as was proposed by Goldman-Rakic and colleagues based on the comparative evidence at the time (Romanski, 2004; Wilson et al., 1993), suggest that the two visual stream architecture may also extend into the human PFC. Using a virtual dissection approach, we related these findings to the underlying white matter association bundles that form the main frontoparietal and frontotemporal connections (Barrett et al., 2020; Thiebaut de Schotten et al., 2011; Eichert et al., 2019). By estimating the cortical projections of the SLF to the FEF and IFJ, we showed that bilaterally, the SLF1 streamlines terminated within the FEF ALE cluster, whereas the SLF3 streamlines terminated within the IFJ ALE cluster. These two branches of the SLF could underlie the connectivity patterns we have reported of the FEF to the medial and superior parietal cortex on the one hand, and the connectivity patterns of the IFJ to the lateral inferior parietal cortex on the other. The SLF2 instead seemed to project to the FEF and IFJ to the same degree, possibly representing a shared pathway mediating their communications with the posterior parietal cortex and its associated functions (Thiebaut de Schotten et al., 2011; Marshall et al., 2015; Parlatini et al., 2017). The connections of the IFJ with the ventral visual stream could be mediated by the AF based on our virtual dissection results. Classically, the AF connects the ventrolateral PFC to areas of the lateral temporal lobe (Fernández-Miranda et al., 2015). A recent study improved our understanding of this bundle (Eichert et al., 2019), revealing that it not only expanded in size but also underwent a major reorganization compared to the macaque (Barrett et al., 2020; but see Becker et al., 2022). According to this study, its surface projections reach the middle and inferior temporal cortex in humans (Eichert et al., 2019), which seems well in line with the IFJ connectivity patterns we reported here.

Our probabilistic tractography pipeline was carefully modeled on the methods reported in Donahue et al. (2016), a study in which the authors were able to assess the relationship between different tractography approaches and retrograde tracer injections taken as a ‘silver standard’ (Sarwar et al., 2021). We relied on the excellent quality and spatial resolution of HCP dMRI data, and we performed all our analyses at the individual subject level, thus preserving as much as possible the detailed anatomy of each brain. We parcellated the cortex according to the MMP1 (Glasser et al., 2016), one of the most comprehensive brain atlases available to date, mapping it to the individual white matter segmentation using surface-based methods, thus exploiting its high parcel resolution, which is often deteriorated in studies using volumetric techniques (Coalson et al., 2018). Our approach was also designed to include as many anatomically plausible constraints as possible based on our hypothesis to optimize the reconstruction of long-range cortico-cortical connections (Jbabdi et al., 2015) and to mitigate some of the biases and common pitfalls of dMRI tractography (Jeurissen et al., 2019; Sotiropoulos & Zalesky, 2017). First, using a pial termination mask allows for discarding streamlines that would incorrectly propagate between adjacent gyri (Hernandez-Fernandez et al., 2019). Second, we chose to seed the streamlines from the WM/GM boundary as several studies show that it is beneficial in alleviating the gyral bias (Girard et al., 2014; Schilling et al., 2018; Van Essen et al., 2014), as well as in reducing partial volume effects (Cottaar et al., 2021; Kleinnijenhuis et al., 2015) and length biases (Girard et al., 2014). Third, by having the MMP1 cortical targets projected on the white matter surface, we strived to improve the assignment of streamline terminations to its labels (St-Onge et al., 2021; Yeh et al., 2019). Finally, several ‘avoid’ masks, which included the segmentations of subcortical nuclei, the corpus callosum, and the cerebellum, were added to discard invalid streamlines that would have potentially resulted in increasing false positive connections (Hernandez-Fernandez et al., 2019).

Our findings highlight a more general anatomical motif of segregated white matter pathways from the PFC and the posterior visual streams, suggesting that the two visual stream architecture may extend into the PFC, as was first proposed in the macaque model (Romanski, 2004; Wilson et al., 1993; Yeterian et al., 2012; see Martinez-Trujillo, 2022, for a recent review). These findings also provide converging evidence for the functional specialization of the FEF and IFJ in spatial versus non-spatial processing in attention and working memory functions (Bedini & Baldauf, 2021; O’Reilly, 2010; see Bichot et al., 2015, 2019, for comparative evidence). Based on its set of input and output connections, the FEF is optimally placed within the visual processing hierarchy to efficiently bias spatial selection and working memory in posterior visual regions (Felleman & Van Essen, 1991; Liu & Hou, 2013; Yeo et al., 2011). Similarly, the IFJ receives inputs and sends feedback signals to regions across the ventral stream to efficiently bias non-spatial (feature-and object-based) attention and working memory (Baldauf & Desimone, 2014; Liu & Hou, 2013; Zanto et al., 2011; Zhang et al., 2018). These multimodal connectivity patterns suggest that the attention networks may be organized in a domain-specific manner. In the spatial domain, it is well established that spatial information is encoded by each region at the neural population level forming topographic maps along the visual hierarchy, culminating in the PFC where they overlap with the FEF (Mackey et al., 2017; Wang et al., 2015). Corresponding topographic representations of the visual field are structurally and functionally wired across adjacent regions in the visual, parietal, and ventral cortex (Finzi et al., 2021; Greenberg et al., 2012; Heinzle et al., 2011; Mohavedian Attar et al., 2020). Recently, preferential topographic patterns of connectivity (both structural and functional, resting-state and task-based) were also uncovered between frontal regions and V1 (Griffis et al., 2015; Knapen, 2021; Sims et al., 2021; Yeo et al., 2011). These patterns of preferential connectivity may provide a computationally efficient architecture for top-down spatial selection (Jbabdi et al., 2013b). In the case of non-spatial domains, it is arguably more difficult to pinpoint an architecture of this kind. Comparative studies have, however, begun to elucidate the organization of the ventrolateral PFC in the form of feature and object-selective cortical patches (Haile et al., 2019; Tsao et al., 2008). We suggest that, in principle, this form of organization could provide a similar basis for preferential topographical connectivity across different processing domains (Xu et al., 2022). Our study ties into this framework by highlighting the white matter pathways associated with different forms of top-down control and therefore opens the possibility of fractionating the attention networks based on the representational content encoded (Bedini & Baldauf, 2021; Liu & Hou, 2013; Osher et al., 2019; Parlatini et al., 2017; Rajan et al., 2021; Soyuhos & Baldauf, 2023).

## 5. CONCLUSION

Here, by combining an ALE fMRI meta-analysis with surface-based probabilistic tractography, we uncovered the structural connectivity of the FEF and the IFJ with the dorsal and ventral visual streams. Our approach, informed by the MMP1 parcellation and leveraging the individual subject level analysis of a large sample size from the HCP dataset, allowed us to provide evidence supporting the view that the two visual streams essentially extend into the human PFC. Our study suggests that the functions of the FEF and IFJ, especially concerning spatial and non-spatial selection (Bedini & Baldauf, 2021), are ultimately grounded in their structural connectivity fingerprints (Osher et al., 2016; Passingham & Lau, 2022; Saygin et al., 2012).

## CRediT AUTHOR CONTRIBUTION STATEMENT

**Marco Bedini:** Conceptualization, Data Curation, Formal Analysis, Funding Acquisition, Investigation, Methodology, Project Administration, Resources, Software, Validation, Visualization, Writing – Original Draft, Writing – Review & Editing; **Emanuele Olivetti:** Methodology, Supervision; **Paolo Avesani:** Methodology, Writing – Review & Editing, Supervision; **Daniel Baldauf:** Conceptualization, Funding Acquisition, Methodology, Writing – Review & Editing, Supervision.

## DATA AVAILABILITY

The structural and diffusion MRI data released by the HCP were downloaded from: https://db.humanconnectome.org. The HCP pipelines can be found at: https://github.com/Washington-University/HCPpipelines. The command-line tools from the FSL FDT toolbox are available at: https://fsl.fmrib.ox.ac.uk/fsl/fslwiki/FSL. The binaries of probtrackX GPU implementation were downloaded from: https://users.fmrib.ox.ac.uk/∼moisesf/Probtrackx_GPU/Installation.html. The code for mapping the MMP1 atlas to the native structural space can be found at: https://figshare.com/articles/dataset/HCP-MMP1_0_volumetric_NIfTI_masks_in_native_stru <u>ctural_space/4249400</u>. Data will be made available on request.

## ACKNOWLEDGMENTS

MB was supported by a doctoral scholarship from the University of Trento. MB would like to thank Lisa Novello for advice on dMRI methods, Gabriele Amorosino for help configuring the NVIDIA GPU drivers on Ubuntu 18.04 LTS and for the discussions on image registration, and Giorgio Marinato for help with shell scripting. Finally, MB is also grateful to the Language, Interaction and Computation Laboratory (CLIC) lab at CIMeC for granting him access to their computing resources. Data were provided in part by the Human Connectome Project, WU-Minn Consortium (Principal Investigators: David Van Essen and Kamil Ugurbil; 1U54MH091657) funded by the 16 NIH Institutes and Centers that support the NIH Blueprint for Neuroscience Research; and by the McDonnell Center for Systems Neuroscience at Washington University.

## Footnotes

1 Present address: Institut de Neurosciences de la Timone, Aix-Marseille University, Faculté de Médecine 27, Boulevard Jean Moulin, 13005 Marseille, France

2 We didn’t normalize the values according to the size of the target region. Although our target regions vary in size and in how they relate to the cortical morphology (i.e., the gyral patterns), and this could partly explain the difference observed in their streamline counts, our focus was on contrasting the connectivity likelihood with each target between the ipsilateral FEF and IFJ seeds. Therefore, although we recognize this factor may play a role in shifting the connectivity likelihoods upward in the case of larger cortical regions, it should have a negligible impact on our subsequent analyses (see Rosen & Halgren, 2021, for analogous considerations on normalization strategies).

