## Supplementary material for "Surface-based Tractography uncovers ‘What’ and ‘Where’ Pathways in Prefrontal Cortex": Suppl. Info

### SUPPLEMENTARY INFORMATION

#### 1. Sample selection criteria

Thanks to the access to restricted demographic information, we retrieved a subset of the HCP data by filtering the initial sample (n = 95) according to the following criteria (see Figure S1 for a schematic): first, we discarded subjects who had documented quality control issues (n = 4); next, we identified all the subjects with family relationships (i.e., twins), and finally we randomly selected a single twin while all the related twins were discarded from the sample. This led to the inclusion of 36 subjects with no family relationship. Finally, we added to this sample all the unrelated subjects from the release who also had resting-state MEG data available (n = 20), since the parallel analysis of the latter modality is part of ongoing work from our lab.

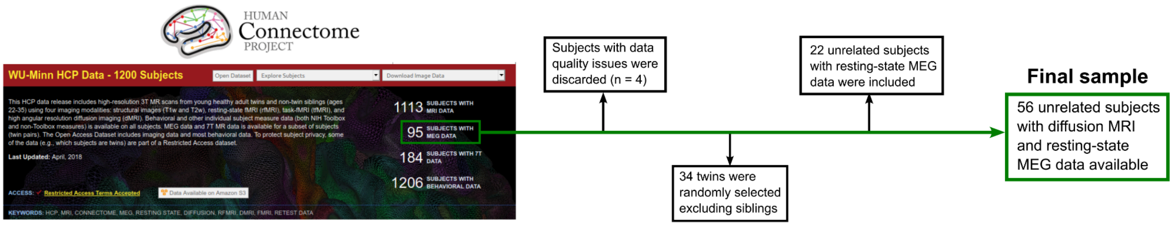


**FIGURE S1** Schematic of the sample selection procedure from the HCP-MEG dataset

[1] Present address: Institut de Neurosciences de la Timone, Aix-Marseille University, Faculté de Médecine 27, Boulevard Jean Moulin, 13005 Marseille, France

### 2. Activation likelihood estimation fMRI meta-analysis method

In the FEF subsample, we included all the tasks that were analyzed in our previous study (i.e., prosaccades > fixation, antisaccades > fixation, and antisaccades > prosaccades, valid > neutral/invalid trials), with the addition of more complex oculomotor contrasts (for examples, see Jarvstad & Gilchrist, 2019, and Thakkar et al., 2014). In contrast, in the IFJ subsample, we included all the studies presented in our previous study except for Meyyappan et al. (2021). Both subsamples included 21 eligible studies from which a single contrast of interest per experiment was used as input for GingerALE. In total, there were 25 experiments in the FEF sample and 22 in the IFJ sample. Thus, even though our samples were lower than the analysis presented by Bedini et al. (2023), this analysis was still adequately powered according to the recommendations by Eickhoff et al. (2016), who recommended 17-20 experiments as a lower boundary. Tables S1 and S2 summarize the information on each included study.

| Study | N | Age | Paradigm | Contrast | Design | Eye tracker | Results reported for | N° of foci | Multiple FEF foci |
|---|---|---|---|---|---|---|---|---|---|
| Amiez and Petrides (2018) | 13 | 22.6 ± 2.8 | Functional localizer | Prosaccades > Fixation | Blocked | N | Single foci | 2 | N |
| Cameron et al. (2015) | 15 | 20.7 ± 1.7 | Functional localizer | Antisaccades & prosaccades > Fixation | *Event-related* | Y | ROIs only | 8 | N |
| Christophel et al. (2018) | 22 | 24.4 ± 0.83 | Functional localizer | Vertical & horizontal instructed saccades > Fixation | *Event-related* | N | Single foci | 2 | N |
| Duecker et al. (2013) | 20 | 19-28 | Functional localizer | Prosaccades > Fixation | Blocked | N | Single foci | 2 | N |
| Fernandez-Ruiz et al. (2018) | 25 | 21.7 ± 1.9 | Functional localizer | Correct prosaccades & antisaccades > Fixation | *Event-related* | Y | Whole-brain | 18 | N |
| Ford et al. (2005) | 10 | 28 | Oculomotor task | Late preparatory period comparison: Antisaccades > Prosaccades | *Event-related* | Y | Whole-brain | 8 | N |
| Furlan et al. (2016) | 6 | NA | Functional localizer | Execution of Antisaccades & Prosaccades > Fixation | *Event-related* | Y | ROIs only | 6 | N |
| Guo et al. (2012) | 12 | 19-31 | Functional localizer | Prosaccades > Fixation | Blocked | Y | ROIs only | 6 | N |
| Gurel et al. (2018) | 16 | 23.2 | Functional localizer | Prosaccades > Fixation | Blocked | Y | Single foci | 2 | N |
| Jamadar et al. (2015) | 23 | 25.8; 18-43 | Functional localizer | Antisaccades > Fixation | *Event-related* | Y | Whole-brain | 47 | N |
| Jarvstad and Gilchrist (2019) | 23 | NA | Functional localizer | Prosaccades in inhibition trials > Prosaccades in selection trials | Blocked | Y | Whole-brain | 4 | N |
| Kastner et al. (2007) | 4 | 20-36 | Functional localizer | Prosaccades > Fixation | Blocked | Y | ROIs only | 5 | N |
| Krebs et al. (2010) | 16 | 25 ± 4.6 | Oculomotor task | Conditional saccades > Fixation | *Event-related* | Y | Whole-brain | 15 | Y |
| Levy et al. (2007) | 8 | 24-43 | Functional localizer | Prosaccades > Fixation | Blocked | N | ROIs only | 5 | N |
| Manoach et al. (2007) | 21 | 34.2 ± 12.6 | Functional localizer | Antisaccade > Prosaccade | *Event-related* | Y | Whole-brain | 19 | N |
| Schon et al. (2008) | 17 | 21.29 ± 3.72 | Functional localizer | Prosaccades > Fixation | Blocked | N | Whole-brain | 19 | Y |
| Tamber-Rosenau et al. (2018) | 10 | 28.5 ± 3.3 | Functional localizer | Prosaccades > Fixation | Blocked | Y | ROIs only | 6 | N |
| Thakkar et al. (2014) | 37 | 29.3; 19-48 | Oculomotor task | Compensated > No-step saccades | *Event-related* | Y | Whole-brain | 39 | Y |
| Tibber et al. (2010) | 16 | 21-40 | Functional localizer | Endogenous & exogenous prosaccades > Fixation | Blocked | Y | ROIs only | 9 | N |
| Van Pelt et al. (2010) | 18 | 20-37 | Functional localizer | Prosaccades in darkness > Fixation | *Event-related* | Y | ROIs only | 7 | N |
| Vossel et al. (2015) | 16 | 24.9; 19-31 | Functional localizer | Cued prosaccades > Fixation | *Event-related* | Y | Whole-brain | 11 | N |

**TABLE S1** Studies included in the FEF sample

| Study | N | Age | Paradigm | Contrast | Design | Results reported for | N° of foci |
|---|---|---|---|---|---|---|---|
| Asplund et al. (2010) | 30 | NA | RSVP / Oddball paradigm | Surprise > Search trials | *Event-related* | Whole-brain | 4 |
| Baldauf and Desimone (2014) | 12 | 23-37 | Object-based attention paradigm | Attend face & attend house blocks > Passive view | Blocked | Single foci | 2 |
| Bollinger et al. (2010) | 18 | 23.4 ± 3.06; 18-28 | Working memory paradigm (n-back) | Functional connectivity with FFA: Stimulus known > Passive view + stimulus unknown | *Event-related* | Whole-brain | 17 |
| Cole and Schneider (2007) | 9 | 19-42 | Working memory paradigm | Target switching trials target non-occluded > Non-switching trials target non-occluded | Mixed blocked/*Event-related* | Whole-brain | 12 |
| Derrfuss et al. (2012) | 12 | 25.3 ± 2.4; 22-31 | Stroop paradigm | Incongruent > Congruent trials | *Event-related* | ROIs only | 2 |
| Han and Marois (2013) | 22 | 19–35 | RSVP / Oddball paradigm | Discontinuous > Continuous & continuous-hard conditions | *Event-related* | Whole-brain | 12 |
| Han and Marois (2014) | 14 | 20-32 | RSVP / Oddball paradigm | Target > Distractor trials | *Event-related* | Whole-brain | 11 |
| Han and Marois (2014) | 6 | 19-35 | RSVP / Oddball paradigm | Target > Distractor trials | *Event-related* | Whole-brain | 11 |
| Han et al. (2018) | 20 | 22-33 | RSVP / Oddball paradigm | Oddball trials > Search trials | *Event-related* | ROIs only | 4 |
| Henseler et al. (2011) | 27 | 24.56 ± 2.53 | Covert attention / Working memory paradigm | Internal > External attending (position task) | Blocked | Whole-brain | 15 |
| Lin et al. (2019) | 18 | 27.4 ± 6.6 | Working memory paradigm (n-back) | Functional connectivity face cue > scene cue | *Event-related* | Whole-brain | 9 |
| Meyyappan et al. (2022) | 20 | 24.65 ± 2.8 | Endogenous cueing paradigm | Spatial and feature-based task cue-evoked activity | *Event-related* | Whole-brain | 23 |
| Sreenivasan et al. (2014) | 16 | 22; 18-32 | Working memory paradigm | Memory > Rotation discrimination trials | Blocked | Whole-brain | 13 |
| Stelzel et al. (2010) | 48 | F: 22 ± 1.99; M: 22.6 ± 1.99 | Task-switching paradigm | Task switch > Repetition trials | *Event-related* | ROIs only | 2 |
| Todd et al. (2011) | 18 | 18-31 | Working memory paradigm (n-back) | Encoding period > No-event fixation trials | *Event-related* | Whole-brain | 27 |
| Wills et al. (2017) | 22 | 18-30 | Contingent capture paradigm | Salient target > Baseline | *Event-related* | ROIs only | 13 |
| Yin et al. (2017) | 26 | 21.3; 21-25 | Task-switching paradigm | Task switch > Repetition trials | *Event-related* | Whole-brain | 15 |
| Zanto et al. (2010) | 13 | 25; 20-31 | Working memory paradigm (n-back) | Functional connectivity V5: Attend > Ignore motion | *Event-related* | Whole-brain | 8 |
| Zanto et al. (2011) | 20 | 24.25; 18-31 | Working memory paradigm (n-back) | Functional connectivity V5: Attend > Ignore motion | *Event-related* | Whole-brain | 16 |
| Zanto et al. (2014) | 20 | 25.7 | Working memory paradigm (n-back) | Functional connectivity V5: Attend > Ignore motion | *Event-related* | ROIs only | 5 |
| Zhang et al. (2018) | 19 | 19-26 | Feature-based attention paradigm | Stimulus block > Baseline | Blocked | ROIs only | 8 |
| Zhao et al. (2020) | 13 | 21.54 ± 1.99 | Working memory paradigm | High working memory load > Low working memory load | *Event-related* | Whole-brain | 7 |

**TABLE S2** Studies included in the IFJ sample

### 2.2 Activation likelihood estimation fMRI meta-analysis results and discussion

We modeled the uncertainty associated with their localization across several hundreds of subjects (378 for the FEF sample, and 423 for the IFJ sample). We found the highest spatial convergence of activations at the junction of the superior frontal sulcus with the superior precentral sulcus for the FEF, and the junction of the inferior frontal sulcus with the inferior precentral sulcus for the IFJ (see Bedini et al., 2023, for an extension of these results). Given the slight differences in the meta-analysis samples compared to our previous study, we obtained generally similar but not completely identical results in the localization of the ALE peaks. The coordinates of the ALE peaks generally matched but were slightly shifted in the case of RH FEF (here LH FEF: -28, -6, 54, and RH FEF: 32, -4, 52, whereas in Bedini et al. (2023) they were LH FEF: -28, -6, 54, and RH FEF: 30, -6, 50), and LH IFJ (LH IFJ: -42, 8, 26, and RH IFJ: 46, 12, 28, whereas in Bedini et al. (2023) they were LH IFJ: -42, 6, 30, and RH IFJ: 46, 12, 28), although they remained identical for the LH FEF and the RH IFJ. To the extent that our tractography approach allowed, these differences should only have minor effects on the results presented in this study, as our seeding approach included the updated ALE peak coordinates corresponding to the results from our previous publication (Bedini et al., 2023).

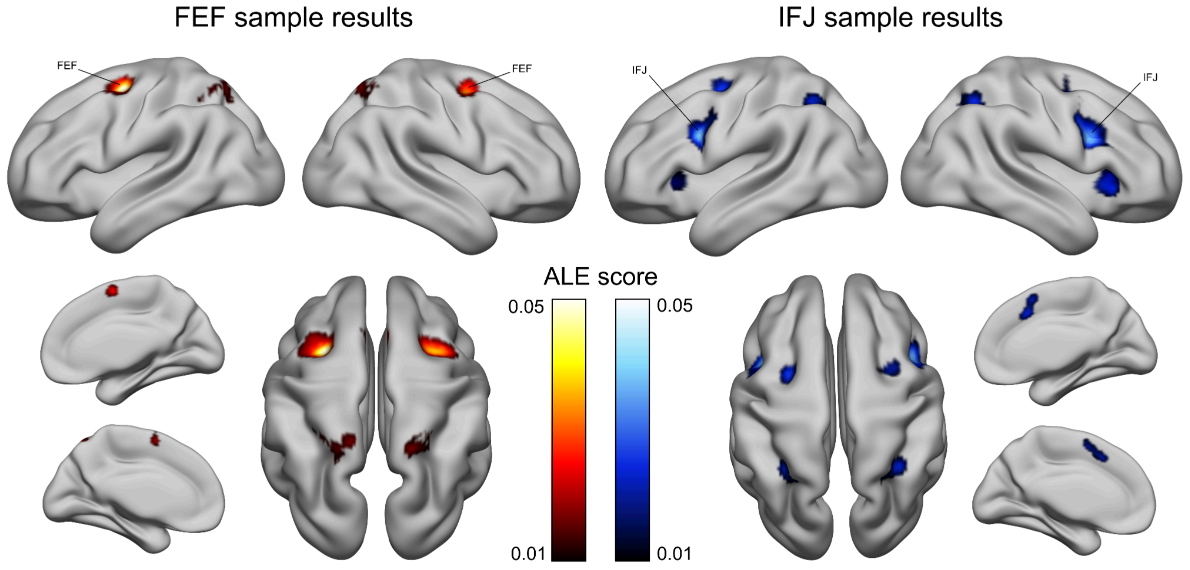


**FIGURE S2** Whole-brain ALE results

### 3. Selection of retinotopic visual regions from the MMP1 and exclusion criteria

To assess whether a parcel could be considered topographically organized, we compared the MMP1 with the atlas by Wang et al. (2015) by overlaying them on FSaverage (Fischl et al., 1999). In particular, we used the maximum probability maps (MPM) from Wang's atlas and hypothesized a correspondence between their regions. Even though these studies relied on different data sources, it is widely assumed that the different cortical features (retinotopy, myelin content, resting-state functional connectivity) have a high degree of alignment (Glasser et al., 2016), especially early in the visual hierarchy.

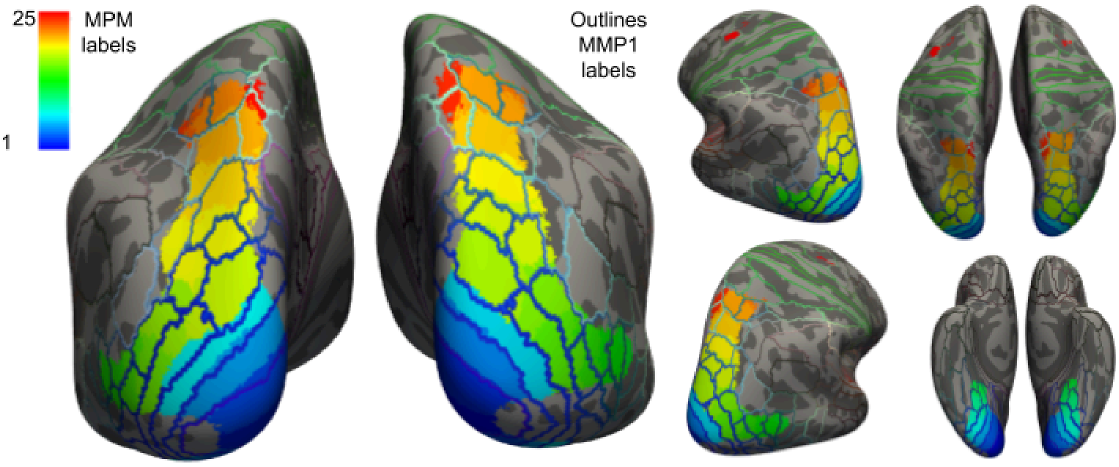


**FIGURE S3** The maximum probability maps (MPM) from Wang et al. (2015) were overlaid on a version of the MMP1 (Glasser et al., 2016) mapped onto the FSaverage surface

Finally, we excluded from our visual streams' definition parcels near the medial wall, and parcels from the superior temporal lobe and the temporo-parietal junction were also excluded since they are likely characterized by distinctive connectional profiles (Kravitz et al., 2013; Mars et al., 2011). Crucially, we also decided not to include early visual regions in our probabilistic tractography analysis, for several reasons. Even though prominent models suggest for example reliable functional interactions between FEF and V4, especially in the macaque model (Moore & Armstrong, 2003), it is unclear whether similar effects are direct or indirect in humans (Kastner & Ungerleider, 2000). Furthermore, there is also converging fMRI and EEG evidence of interactions between IFJ and V4 (Zanto et al., 2011; Zhang et al., 2018), which however again leaves open the question of whether these effects are direct or indirect. Another motivation relates to the fact that while all the early visual regions are topographically organized (Wang et al., 2015), some display spatial and feature selectivity, hence potentially constituting a zone of graded influence from either the FEF or the IFJ. In summary, based on these arguments, we decided to exclude early visual cortex regions from our analyses.

### 4. Surface-based probabilistic tractography results

Full results of the contrasts between contrast between the FEF and IFJ normalized streamline counts for the left hemisphere dorsal stream targets (Figure S4), left hemisphere ventral stream targets (Figure S5), right hemisphere dorsal stream targets (Figure S6) and right hemisphere ventral stream targets (Figure S7). A value of −25 is set as an arbitrary $\log_2$ connectivity likelihood equal to 0 for visualization convenience in all the figures.

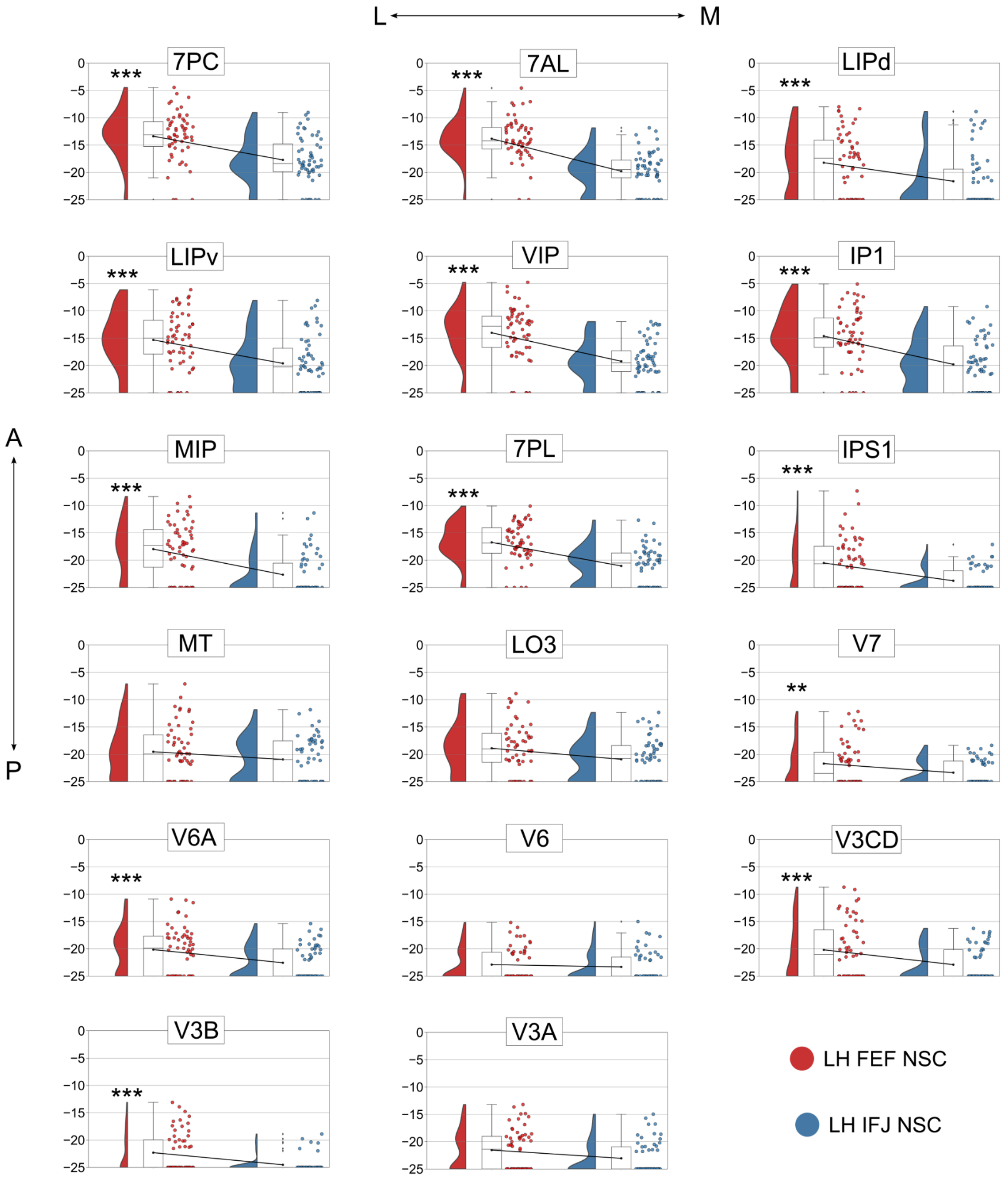


**FIGURE S4** Plots of the $\log_2$-scaled normalized streamline counts (NSC) of the FEF and IFJ to all the cortical targets in the left hemisphere dorsal visual stream

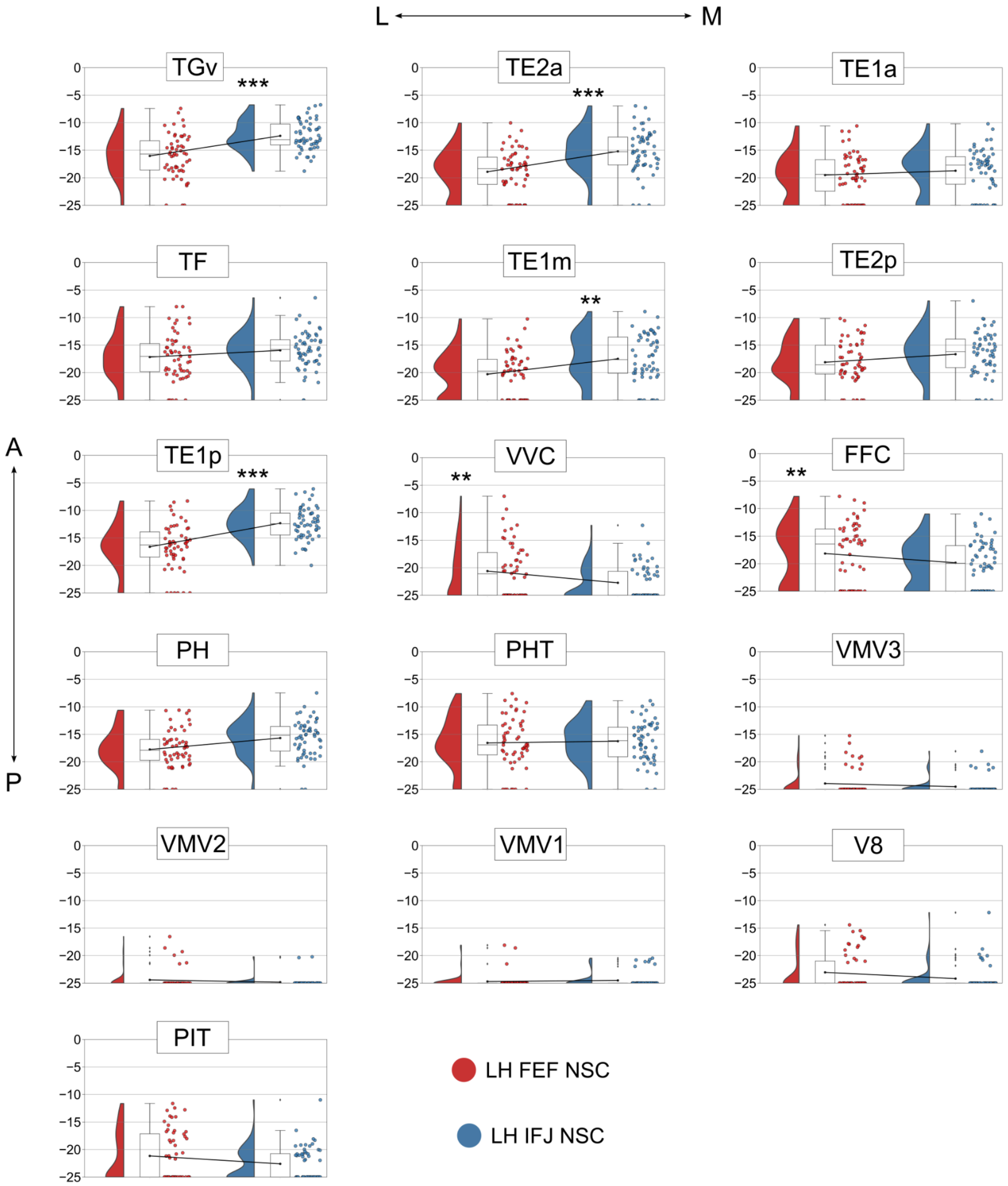


**FIGURE S5** Plots of the $log_2$-scaled normalized streamline counts (NSC) of the FEF and IFJ to all the cortical targets in the left hemisphere ventral visual stream

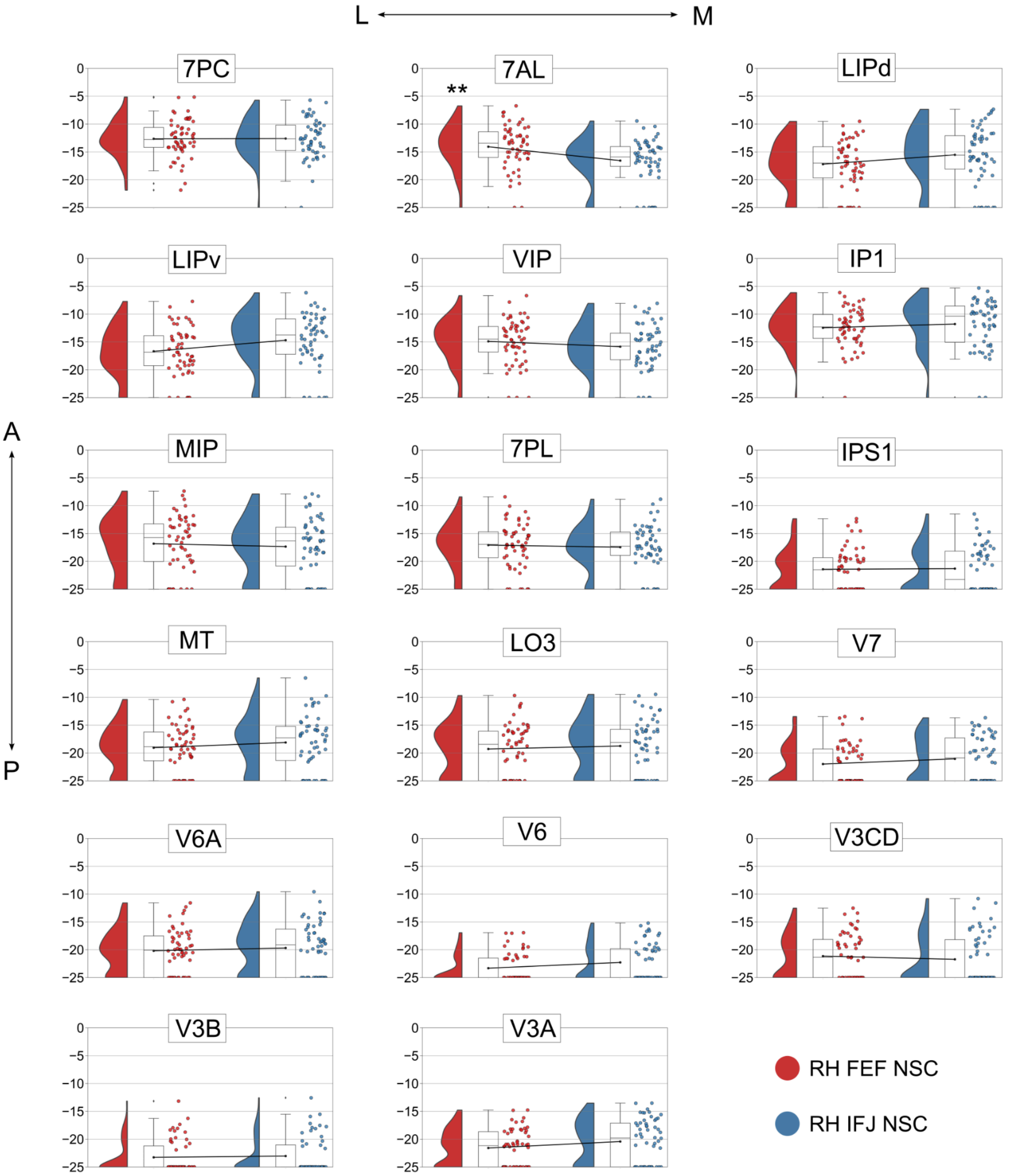


**FIGURE S6** Plots of the $log_2$-scaled normalized streamline counts (NSC) of the FEF and IFJ to all the cortical targets in the right hemisphere dorsal visual stream

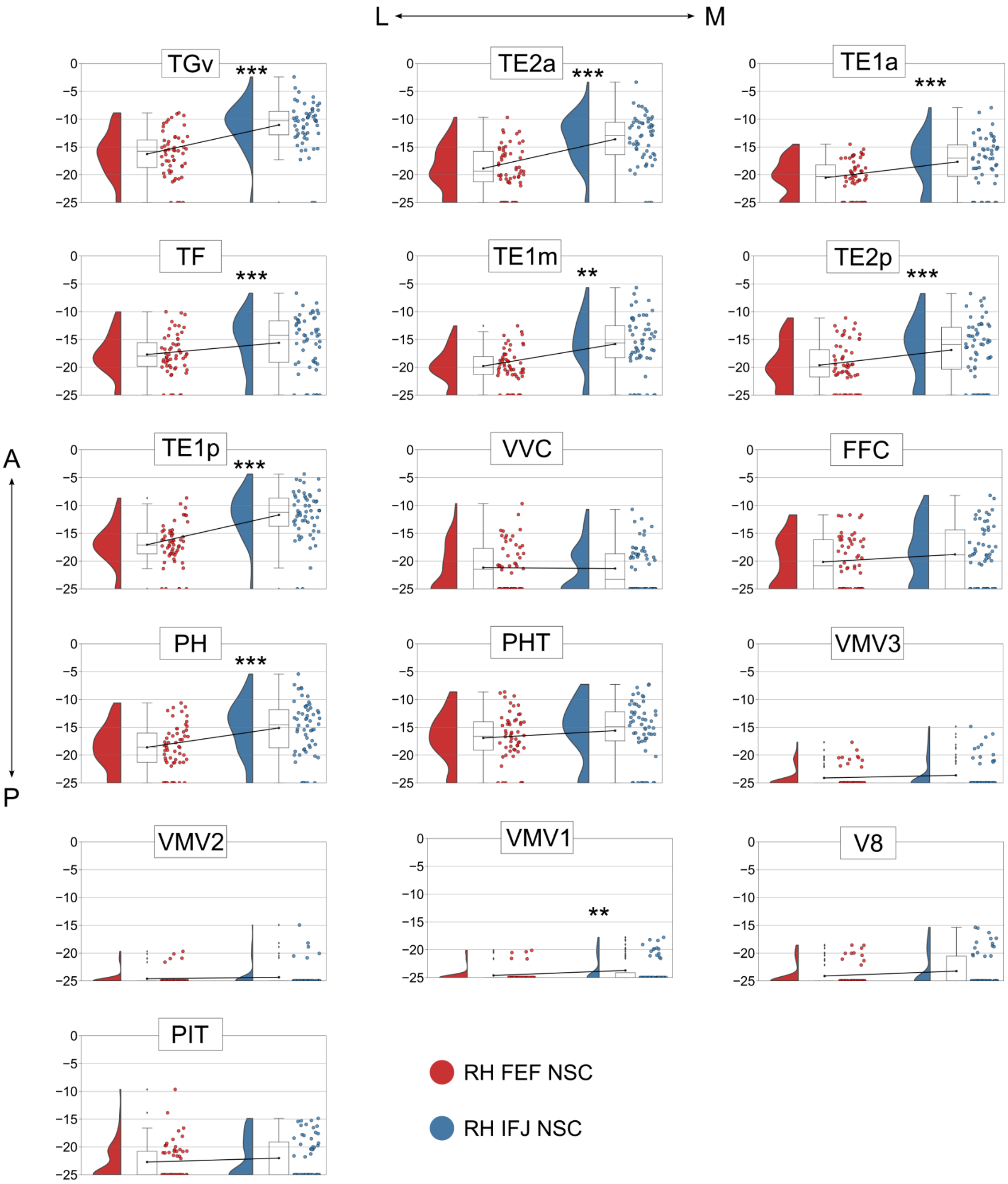


**FIGURE S7** Plots of the $\log_2$-scaled normalized streamline counts (NSC) of the FEF and IFJ to all the cortical targets in the right hemisphere ventral visual stream

### 4.1 Limitations

Even though we relied on state-of-the-art FSL tools and validated pipelines for tractography (Donahue et al., 2016) and applied the a priori anatomical constraints mentioned, the presence of false positive connections is a limitation that could still affect some of our results (Maier-Hein et al., 2017). It is indeed likely that densely sampling the fiber orientations inferred with bedpostx using 50000 streamlines per vertex to improve the sensitivity of our analysis may have also led to inadvertently increasing the number of non-plausible streamlines. Therefore, we acknowledge that using a computational technique to filter out these streamlines would be beneficial (Pestilli et al., 2014; Sotiropoulos & Zalesky, 2017). Another potentially critical aspect of our analysis concerns our seeding strategy. We only tracked from the FEF and IFJ to the targets, and not symmetrically, from these seeds to targets and vice versa (as in Rosen & Halgren, 2021). Although it seems reasonable to hypothesize that the connections we reported here are largely reciprocal, and comprise afferent and efferent connections (and hence, that we could potentially replicate these results by tracking from the posterior visual streams), our method doesn’t technically allow us to distinguish between these possibilities (Jbabdi & Johansen-Berg, 2011).

### 5.1 Method for the analysis of bundle asymmetries

Analyzing bundle asymmetries gave us a way to indirectly carry out a sanity check on our bundle segmentation results, as we expected to find results largely in line with the literature on bundle lateralization. In addition, this also allowed us to assess potential differences between the two hemispheres that may provide complementary evidence to our surface-based probabilistic tractography analysis. The bundles of interest were normalized by their streamline count, binarized at 0.5%, and a lateralization index (L) was computed using the formula (where V = volume; as in de Schotten et al., 2011, and Warrington et al., 2020):

$$L = \frac{V\,RH - V\,LH}{V\,RH + V\,LH}$$

The normality assumption was checked for all bundles with the Shapiro-Wilk test, and we assessed bundle lateralization using one-sample *t*-tests ($p = 0.01$).

### 5.2 Results of the analysis of bundle asymmetries

Our results show that the AF (t(23), p = 0.02, d = −0.512) and the UF were left-lateralized (t(23), p < 0.001, d = −1.198). In contrast,, the SLF2 was right-lateralized (t(23), p = 0.015, d = 0.538).

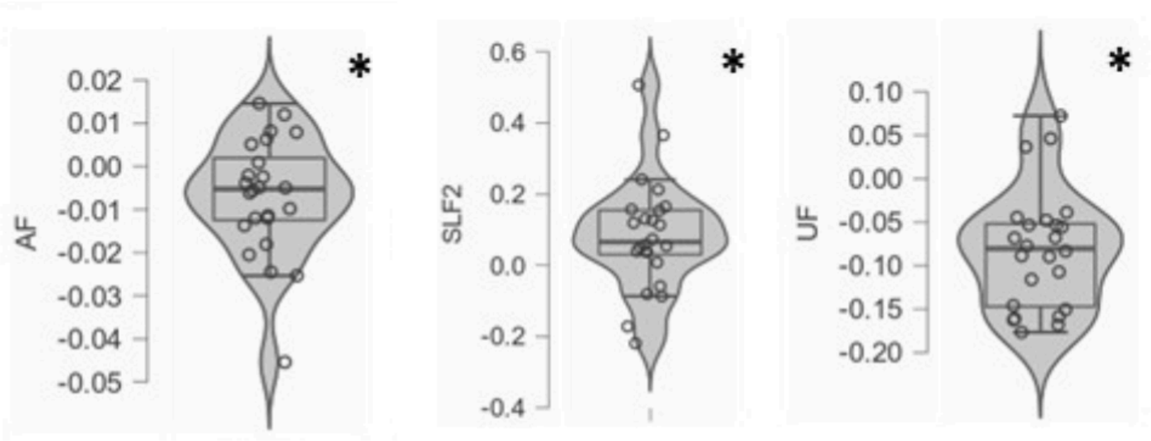


**FIGURE S8** Bundles that showed significant hemispheric asymmetries

### 5.3 Discussion of the results

Interestingly, the fact that the SLF2 was right-lateralized, but not the SLF3, is somewhat in contrast with previous studies (Marshall et al., 2015; Thiebaut de Schotten et al., 2011; Warrington et al., 2020), where only the latter showed that pattern. This could be due to several reasons: first, even though the XTRACT method (Warrington et al., 2020) is based on these previous studies for what concerns the definition of the ROIs for the virtual dissection, it is an automatic method and hence manual intervention may be helpful in some instances to refine the bundle segmentations. Secondly, the algorithm it leverages (probtrackx) is also different from the other studies cited, which may lead to discordant results. Third, as we removed any family relationships from the sample to make it more representative of the general population, our selection strategy increased interindividual variability in the sample. This factor is known to play a role in SLF lateralization (Bartolomeo & Seidel Malkinson, 2019). Warrington et al. (2020) reported that the SLF2 was left-lateralized in the HCP dataset, and in contrast, it was right-lateralized in the UK Biobank dataset.

Our results on the left lateralization of the AF and UF are on the other hand very much in line with the literature (for the AF, see Eichert et al., 2019, and Fernández-Miranda et al., 2015). These results were also replicated by Warrington et al. (2020) and were consistent in the HCP and UK Biobank datasets.
